# Gut microbiome diversity detected by high-coverage 16S and shotgun sequencing of matched stool and colon biopsy samples

**DOI:** 10.1101/742635

**Authors:** Joan Mas-Lloret, Mireia Obón-Santacana, Gemma Ibáñez-Sanz, Elisabet Guinó, Miguel L Pato, Francisco Rodriguez-Moranta, Alfredo Mata, Ana García-Rodríguez, Victor Moreno, Ville Nikolai Pimenoff

## Abstract

The gut microbiome has a fundamental role in human health and disease. However, studying the complex structure and function of the gut microbiome using next generation sequencing is challenging and prone to reproducibility problems due to the heterogeneity of sample sets. Here, we obtained cross-sectional colon biopsies and faecal samples from nine participants in our COLSCREEN study and sequenced them in high coverage using Illumina pair-end shotgun (for faecal samples) and IonTorrent 16S (for paired feces and colon biopsies) technologies. The metagenomes consisted of between 47 and 92 million reads per sample and the targeted sequencing covered more than 300K reads per sample across seven hypervariable regions of the 16S gene. Our data is freely available and coupled with code for the presented metagenomic analysis using up-to-date bioinformatics algorithms. These results will add up to the informed insights into designing comprehensive microbiome analysis and also provide data for further testing for unambiguous gut microbiome analysis.

## Background & Summary

The gut microbiome is highly dynamic and variable between individuals, and is continuously influenced by factors such as individual’s diet and lifestyle[1, 2], as well as host genetics[3]. Next generation sequencing (NGS) has greatly enhanced our understanding of the human microbiome, as these techniques allow to investigate abundance and diversity of bacteria in a culture-independent manner. Recent developments in bioinformatics have permitted the identification of thousands of novel bacterial and archaeal species and strains identified in human and non-human environments through metagenome assembly[4, 5, 6]. For colorectal cancer (CRC), recent large-scale studies have revealed specific faecal microbial signatures associated with malignant gut transformations, although the causal role of gut bacterial taxa in CRC development is still unclear[7, 8].

The 16S small subunit ribosomal gene is highly conserved between bacteria and archaea, and thus has been extensively used as a marker gene to estimate microbial phylogenies[9]. The 16S rRNA gene contains nine hypervariable regions (V1-V9) with bacterial species-specific variations that are flanked by conserved regions. Hence, the amplification of 16S rRNA hypervariable regions can be used to detect microbial communities in a sample usually up to the genus level[10]. Species-level assignment is also possible if full-length 16S sequences are retrieved[11].

However, conserved regions are not entirely identical across groups of bacteria and archaea, which can have an effect on the PCR amplification step. Notably, among the conserved regions of the 16S gene, central regions are more conserved, suggesting that they are less susceptible to producing bias in PCR amplification[12]. Furthermore, an *in silico* study has shown that the V4-V6 regions perform better at reproducing the full taxonomic distribution of the 16S gene[13]. In another study, a constructed mock sample was sequenced by IonTorrent technology, demonstrating that the V4 region (followed by V2 and V6-V7) was the most consistent for estimating the full bacterial taxonomic distribution of the sample[14]. In addition, other factors such as the actual primer sequence, sequencing technology and the number of PCR cycles may impact on microbiome detection when using 16S sequencing. However, the relative ratios in taxonomic abundance have been shown to be consistent regardless of the experimental strategy used[15].

Beyond 16S sequencing, shotgun metagenomics allows not only taxonomic profiling at species level[16, 17], but also strain-level detection of particular species[18], as well as functional characterization and *de novo* assembly of metagenomes[19]. However, shotgun metagenomics is more expensive than 16S sequencing and may not be feasible when the amount of host DNA is high[20]. Nevertheless, provided sufficient sequencing coverage, taxonomic profiling of shotgun metagenomes is rather robust and mostly depends on the input DNA quality and bioinformatics analysis tools[21]. Indeed, a diverse set of new computational methods and query databases are currently available for comprehensive shotgun metagenomics analysis[22].

Taken together, 16S and shotgun microbiome profiles from the same samples are not entirely the same, but rather represent the relative microbiome composition captured by each methodological approach[23, 24, 25]. In agreement, comparative studies have already revealed that faecal, rectal swab and colon biopsy samples collected from the same individuals usually produce differential microbiome structures although consistent relative taxon ratios and particular core profiles are also detected[26].

In this study, we characterized the gut microbiome signature of nine faecal and colon tissue samples from the same participants. Our data shows a high concordance between different sequencing methods and classification algorithms for the full microbiome on both sample types. However, clear deviations depending on the sample, method, genomic target and depth of sequencing data were also observed, which warrant consideration when conducting large-scale microbiome studies. Therefore, this annotated, high-quality gut microbiome dataset will provide useful insights for designing comprehensive microbiome analysis, and thus, provide data for further testing for unambiguous gut microbiome analysis. Moreover, we provide the full source code for the bioinformatics analysis, available and thoroughly documented on a GitLab repository[27].

## Methods

### Subjects and sampling

The COLSCREEN study is a cross-sectional study that was designed to recruit participants from the Colorectal Cancer Screening Program conducted by the Catalan Institute of Oncology. This program invites men and women aged 50-69 to perform a biennial faecal immunochemical test (FIT, OC-Sensor, Eiken Chemical Co., Japan). Patients with a positive test result (≥20 g Hb/g faeces) are referred for colonoscopy examination. A detailed description of the screening program is provided elsewhere[28, 29]. Exclusion criteria are as follows: gastrointestinal symptoms; family history of hereditary or familial colorectal cancer (2 first-degree relatives with CRC or 1 in whom the disease was diagnosed before the age of 60 years); personal history of CRC, adenomas or inflammatory bowel disease; colonoscopy in the previous five years or a FOBT within the last two years; terminal disease; and severe disabling conditions.

Participants provided written informed consent and underwent a colonoscopy. A week prior to colonoscopy preparation, participants were asked to provide a faecal sample and store it at home at −20 °C. The day of the colonoscopy, participants delivered the faecal sample and a leaflet where they had to report antibiotics, probiotics or laxatives intake in the previous month. Samples under antibiotic or probiotic effect were not analysed. All stool samples were stored in −80°C, while colonic mucosa biopsy samples were performed during the colonoscopy. Four biopsies of normal tissue of each colon segment (4 of ascending colon, 4 of transverse colon, 4 of descending or sigmoid colon, and 4 of rectum) were obtained. If a tumour or a polyp was biopsied or removed, a biopsy was obtained if the endoscopist considered it possible. Subsequently, biopsy samples were immediately transferred to RNAlater (Qiagen) and stored at −80°C.

Colonic lesions were classified according to “European guidelines for quality assurance in CRC”[30]. For the present study, we selected patients with no lesions in the colonoscopy, patients with intermediate-risk lesions (3–4 tubular adenomas measuring <10mm with low-grade dysplasia or as ≥1 adenoma measuring 10-19mm) and with high-risk lesions (≥5 adenomas or ≥1 adenoma measuring ≥20mm). We analysed and compared 18 biological samples (9 faecal samples and 9 colon tissue samples) from 9 participants: n=3 negative colonoscopy, n=3 high-risk lesions, n=3 intermediate-lesions) (Table 1). Our CRC screening programme follows the Public Health laws and the Organic Law on Data Protection. All procedures performed in the study involving data from human participants were in accordance with the ethical standards of the institutional research committee, and with the 1964 Helsinki Declaration and its later amendments or comparable ethical standards. The protocol of the study was approved by the Bellvitge University Hospital Ethics Committee, registry number PR084/16.

**Table 1:** List of samples and DNA concentrations. HRA = high-risk adenoma; IRA = intermediate-risk adenoma; neg = healthy. DNA concentration is presented in ng/*μ*l.

| Sample | Sex | Age | FIT result | Condition | DNA (stool) | DNA (tissue) |
| --- | --- | --- | --- | --- | --- | --- |
| AE1235 | Male | 62 | - | HRA | 17.5 | 4.6 |
| AE1236 | Female | 67 | - | neg | 12 | 7.6 |
| AE1237 | Female | 63 | + | HRA | 12 | 7.9 |
| AE1238 | Male | 61 | - | IRA | 28 | 7.7 |
| AE1239 | Female | 54 | + | neg | 14.9 | 5.7 |
| AE1240 | Male | 63 | - | neg | 10.2 | 4.6 |
| AE1241 | Female | 68 | + | IRA | 15.7 | 3.2 |
| AE1242 | Female | 67 | + | IRA | 18.5 | 6.8 |
| AE1243 | Female | 55 | + | HRA | 6.7 | 4.3 |

### DNA extraction and sequencing

Total faecal DNA was extracted using the NucleoSpin Soil kit (Macherey-Nagel, Duren, Germany) with a protocol involving a repeated bead beating step in the sample lysis for complete bacterial DNA extraction. Total DNA from the snap-frozen gut epithelial biopsy samples was extracted using an in-house developed proteinase K (final concentration 0.1ug/uL) extraction protocol with a repeated bead beating step in the sample lysis. All extracted DNA samples were quantified using Qubit dsDNA kit (Thermo Fisher Scientific, Massachusetts, USA) and Nanodrop (Thermo Fisher Scientific, Massachusetts, USA) for sufficient quantity and quality of input DNA for shotgun and 16S sequencing. DNA yields from the extraction protocols are shown in Table 1.

Metagenomics sequencing libraries were prepared with at least 6.7 ng/uL of total DNA using the Nextera XT DNA sample Prep Kit (Illumina, San Diego, USA) with an equimolar pool of libraries achieved independently based on Agilent High Sensitivity DNA chip (Agilent Technologies, CA, USA) results combined with SybrGreen quantification (Thermo Fisher Scientific, Massachusetts, USA). The indexed libraries were sequenced in one lane of a HiSeq 4000 run in 2×150bp paired-end reads, producing a minimum of 50 million reads/sample at high quality scores (Q30=92.15).

Targeted 16S sequencing libraries were prepared using Ion 16S Metagenomics Kit (Life Technologies, Carlsbad, USA) in combination with Ion Plus Fragment Library kit (Life Technologies, Carlsbad, USA) and loaded on a 530 chip and sequenced using the Ion Torrent S5 system (Life Technologies, Carlsbad, USA). The protocol was designed for microbiome analysis using Ion torrent 510/520/530 Kit-chef template preparation system (Life Technologies, Carlsbad, USA) and included two primer sets that selectively amplified seven hyper-variable regions (V2, V3, V4, V6, V7, V8, V9) of the 16S gene. At least 10 ng of total DNA was used for 16S library preparation and re-amplified using Ion Plus Fragment Library kit for reaching the minimum template concentration. Equimolar pool of libraries were estimated using Agilent High Sensitivity DNA chip (Agilent Technologies, CA, USA). Library preparation and 16S sequencing was performed with the technological infrastructure of the Centre for Omic Sciences (COS).

### Bioinformatics analysis

#### Removal of human reads

Prior to submission of the raw sequence data to the European Nucleotide Archive (ENA), human reads were removed from the metagenome samples in order to follow legal privacy policies. Raw reads were aligned to the human genome (GRCh38) using `Bowtie2` with options `-very-sensitive-local` and `-k 1`. A FASTQ file was then generated from reads which did not align (carrying SAM flag 12) using `Samtools`. These FASTQ files were deposited to the ENA.

#### Shotgun reads quality control

Shotgun samples were quality controlled using `FASTQC`. Accordingly, sequences were deduplicated using `clumpify` from the `BBTools` suite, followed by quality trimming (PHRED > 20) on both ends and adapter removal using `BBDuk`. Read pairs where one read had a length lower than 75 bases were discarded. Results of this quality control pipeline are shown in table 2.

**Table 2:** Output from our quality control pipeline. Numbers indicate the amount of read pairs originally, passing quality control, failing due to duplication or failing due to quality or adapter trimming.

| Sample | Non-human | Clean | Deduplicated | Trimmed |
| --- | --- | --- | --- | --- |
| AE1235 | 27,510,304 | 19,991,742 | 2,042,335 | 5,476,227 |
| AE1236 | 45,050,043 | 29,097,088 | 5,616,498 | 10,336,457 |
| AE1237 | 25,720,634 | 18,745,351 | 2,001,998 | 4,973,285 |
| AE1238 | 34,831,431 | 25,727,431 | 2,708,788 | 6,395,212 |
| AE1239 | 36,353,427 | 25,946,121 | 2,964,013 | 7,443,293 |
| AE1240 | 31,699,249 | 23,225,137 | 2,562,061 | 5,912,051 |
| AE1241 | 34,083,370 | 24,830,987 | 2,739,105 | 6,513,278 |
| AE1242 | 31,592,814 | 23,239,834 | 2,455,849 | 5,897,131 |
| AE1243 | 23,476,326 | 17,887,436 | 1,831,023 | 3,757,867 |

#### Shotgun taxonomic and functional profiling

Pre-processed paired-end shotgun sequences were classified using three different classifiers. `Kraken2` was run against a reference database containing all Ref-Seq bacterial and archaeal genomes (built in May 2019) with a 0.1 confidence threshold. Following classification by `Kraken`, `Bracken` was used to re-estimate bacterial abundances at taxonomic levels from species to phylum using a read length parameter of 150. `MetaPhlAn2` was run using default parameters. `Kaiju` was run against the Progenomes database (built in February 2019) using default parameters. Corresponding taxonomic profiles at family-level are shown in Fig. 1a.

**Figure 1:**
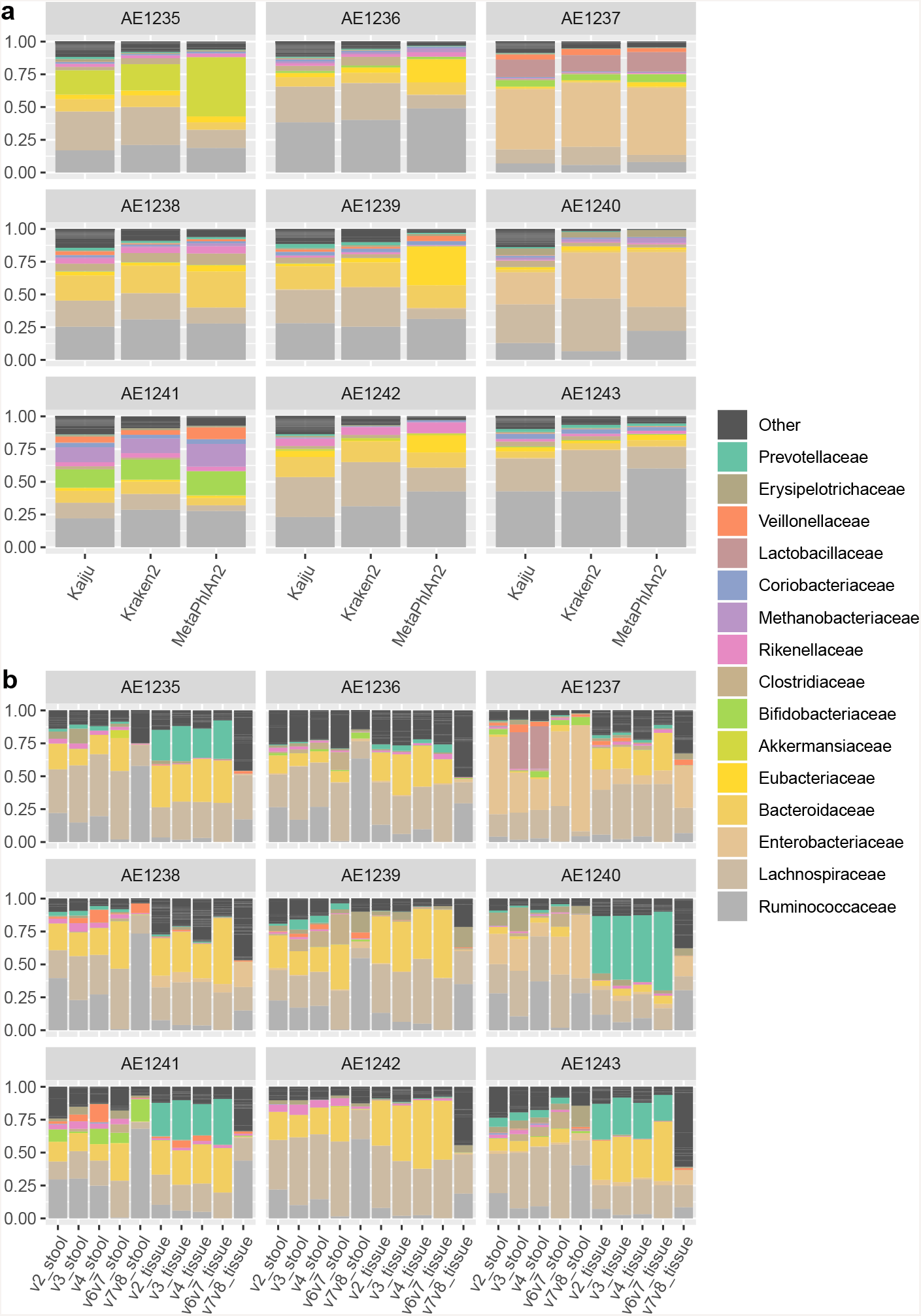
Taxonomic classification of samples at family level. a. Classification of shotgun samples using three different classifiers. b. Classification of 16S sequences, split by region and source material, using DADA2 and IdTaxa.

Functional profiling of the concatenated metagenomic paired-end sequences was performed using the `HUMAnN2` pipeline with default parameters, obtaining gene family (UniRef90), functional groups (KEGG orthogroups) and metabolic pathway (MetaCyc) profiles.

#### De novo assembly

High quality metagenomic reads were assembled using `metaSPADES` with default parameters and binned into putative metagenome assembled genomes (MAGs) using `metaBAT. checkM` was used to check the quality of MAGs and filter them to comply with strict quality requirements (completeness > 90%, contamination < 5%, number of contigs < 300 %, N50 > 20,000). A total of 112 high quality MAGs were assembled from the nine high-coverage metagenomes and assigned a species-level taxonomy using `PhyloPhlAn2`. Assembled species shared by at least two of the nine samples are listed in Table 3.

**Table 3:**
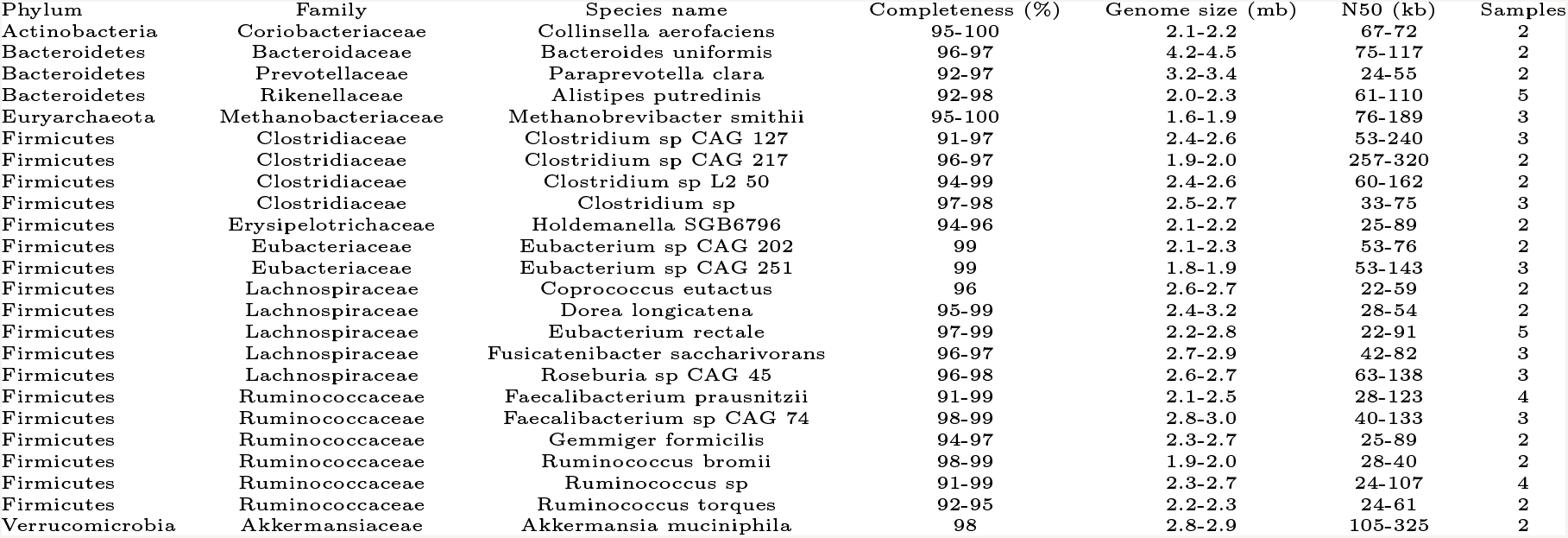
Summary of high quality bins present in at least two samples.

#### Generation of lower coverage pseudo-samples

Pseudo-samples of lower coverage were generated *in silico* using the `reformat` tool from the `BBTools` suite. Five samples were created at 15M, 10M, 5M, 2.5M, 1M, 500K, 100K and 50K read pairs coverage.

Pseudo-samples were then classified using `Kraken2` and `HUMAnN2`. From this classification, Shannon index alpha diversity profiles were computed at the species, genus and phylum level, as well as UniRef90, KO and MetaCyc pathways level using the `R` package `vegan`.

#### Splitting 16S samples by region

An in-house Python program was written in order to separate the 16S sequences according to the variable region(s) in the Ion Torrent 16S dataset. First, we positioned the 16S conserved regions[12] in the E. coli str. K-12 substr. MG1655 16S reference gene (SILVA v.132 Nr99 identifier U00096.4035531.4037072) as well as the corresponding variable region positions[10]. Regions 5 and 7 were truncated to match the reference *E. coli* sequence. Each sequencing read is then assigned into its corresponding variable region by mapping conserved regions into the read taking into account mapping positions.

Analysis of the regions covered in our samples revealed a prevalence of V3, followed by V4, V2, V6-V7 and V7-V8 (Table 4). Each hypervariable region dataset was subsequently extracted from the original FASTQ file with an in-house Python script (code availability).

**Table 4:** Percentage of 16S reads covering each region by sample.

|  |  | Total | V2 | V3 | V4 | V6-V7 | V7-V8 | Other |
| --- | --- | --- | --- | --- | --- | --- | --- | --- |
| Faeces | AE1235 | 739819 | 3.2 | 40.2 | 14.3 | 21.6 | 18.8 | 1.9 |
|  | AE1236 | 450511 | 2.9 | 43.6 | 15.0 | 20.6 | 16.0 | 2.0 |
|  | AE1237 | 767495 | 4.1 | 36.0 | 14.4 | 17.6 | 24.8 | 3.2 |
|  | AE1238 | 740788 | 3.6 | 38.5 | 14.5 | 20.6 | 21.0 | 1.8 |
|  | AE1239 | 997171 | 5.9 | 36.1 | 14.2 | 24.2 | 17.6 | 2.0 |
|  | AE1240 | 458735 | 2.4 | 39.0 | 13.5 | 17.3 | 24.8 | 2.9 |
|  | AE1241 | 590541 | 3.5 | 40.0 | 14.0 | 19.6 | 21.0 | 1.9 |
|  | AE1242 | 467170 | 3.4 | 37.8 | 14.7 | 19.7 | 22.6 | 1.9 |
|  | AE1243 | 386045 | 3.3 | 41.0 | 14.6 | 21.0 | 18.1 | 2.0 |
| Tissue | AE1235 | 321453 | 4.3 | 61.1 | 14.2 | 15.1 | 4.5 | 0.9 |
|  | AE1236 | 621908 | 8.3 | 46.8 | 16.7 | 18.7 | 8.7 | 0.8 |
|  | AE1237 | 726770 | 8.2 | 43.8 | 17.5 | 18.4 | 11.0 | 1.1 |
|  | AE1238 | 735109 | 7.4 | 42.3 | 18.7 | 17.8 | 11.5 | 2.3 |
|  | AE1239 | 577808 | 6.8 | 49.1 | 16.5 | 20.7 | 6.2 | 0.8 |
|  | AE1240 | 601785 | 9.5 | 42.3 | 19.1 | 21.4 | 6.6 | 1.0 |
|  | AE1241 | 649667 | 7.9 | 45.7 | 17.3 | 24.9 | 3.4 | 0.8 |
|  | AE1242 | 589330 | 5.4 | 50.4 | 16.6 | 23.2 | 3.6 | 0.9 |
|  | AE1243 | 447223 | 7.0 | 48.0 | 19.4 | 16.7 | 8.1 | 0.8 |

#### 16S denoising and taxonomic binning

16S sequences were denoised following the standard `DADA2` pipeline with adaptations to fit our single-end read data. For this analysis, reads spanning different regions were introduced into the pipeline as different input files. Taxonomic classification of the high-quality sequences was performed using `IdTaxa` included in the `DECIPHER` package. Taxonomic assignment at family level by region and source material is shown at Fig.1b.

### Statistical analysis

For the statistical analysis of the bacterial abundance data, we employed compositional data analysis methods[31].

Count matrices of the classified taxa were subjected to central log ratio (CLR) transformation after removing low-abundance features and including a pseudo-count. Here, we used the `codaSeq.filter`, `cmultRepl` and `codaSeq.clr` functions from the `CodaSeq` and `zCompositions` packages. Principal components analysis (PCA) biplots were generated from the central log ratios using the `prcomp` function in R.

### Code availability

Software versions for data generation and analysis are listed in Table 5.

**Table 5:** Software versions

| Software | Use | Version |  |
| --- | --- | --- | --- |
| Bowtie2 | Human reads mapping | 2.3.4 | [32] |
| Samtools | Extraction of non-human reads | 1.8 | [33] |
| FASTQC | Reads quality assessment | 0.11.7 | [34] |
| Clumpify | Removal of duplicate reads | 38.26 | [35] |
| BBDuk | Quality and adapter trimming | 38.26 | [35] |
| Kraken | Taxonomic classification of shotgun reads | 2.0.8-beta | [36] |
| Bracken | Re-estimation of taxonomic profiles | 2.2 | [37] |
| MetaPhlAn2 | Taxonomic classification of shotgun reads | 2.7.7 | [38] |
| Kaiju | Taxonomic classification of shotgun reads | 1.6.3 | [39] |
| HUMAnN2 | Functional profiling of shotgun reads | 0.11.1 | [40] |
| metaSPADES | Metagenomic assembly | 3.13.1 | [41] |
| metaBAT | Binning of scaffolds | 2.12.1 | [42] |
| checkM | Bins quality assessment | 1.0.13 | [43] |
| PhyloPhlAn2 | Taxonomic classification of bins | 0.35 | [44] |
| Reformat | Generation of lower coverage samples | 38.26 | [35] |
| DADA2 (R) | Denosing of 16S reads | 1.10.1 | [45] |
| IdTaxa (R) | Taxonomic classification of 16S sequences | 2.10.1 | [46] |
| vegan (R) | Computation of alpha diversity | 2.5.3 | [47] |
| zCompositions (R) | Compositional data analysis | 0.99.3 | [48] |
| CoDaSeq (R) | Compositional data analysis | 1.2.0 | [49] |

Code for sequence quality control and trimming, shotgun and 16S metagenomics profiling and generation of figures in this paper is freely available and thoroughly documented at https://gitlab.com/JoanML/colonbiome-pilot. This repository includes instructions for the analysis and reproduction of the figures on this paper from the publicly available samples, as well as pipelines used for the analysis. This repository is arranged in folders, each containing a README:

- `qc`: Scripts for quality control and preprocessing of samples
- `analysis_shotgun`: Scripts to run softwares for metagenomics analysis
- `regions_16s`: In-house scripts for splitting IonTorrent reads into new FASTQ files
- `analysis_16s`: DADA2 pipeline adapted to this dataset
- `assembly`: Scripts to run the assembly, binning and quality control software
- `figures`: Scripts used to generate the figures in this manuscript

## Data Records

The raw sequence data generated in this work were deposited in European Nucleotide Archives (ENA). Faecal metagenomic sequences are available under accession PRJEB33098. Faecal 16S sequences are available under accession PRJEB33416 and tissue 16S sequences are available under accession PRJEB33417. Human sequences were removed from whole shotgun samples as previously described prior to the ENA submission.

## Technical Validation

Prior to analysis, shotgun sequencing reads were subject to quality and adaptor trimming as previously described. Moreover, reads were deduplicated to avoid compositional biases caused by PCR duplicates. Quality control and denoising of 16S reads was performed within the `DADA2` denoising pipeline and not as an independent data processing step.

In order to validate our pipeline for 16S variable region assignment of reads, we used sequences that were assigned to a species by the `assignSpecies` function in `DADA2`, which searches for unambiguous full-sequence matches in the SILVA database. These sequences were aligned to a randomly selected full 16S gene from the SILVA database (version 132). As shown in Table 6, sequences mapped in expected positions within the 16S gene in concondance to the variable region assigned by our pipeline. This indicates that our pipeline was able to assign variable regions properly.

**Table 6:** Region(s) covered by reads and the position they mapped to within the 16S gene. SILVA IDs: *B. fragilis*: FQ312004.3243020.3244552; *B. vulgaris*: CP000139.2183533.2185042; *F. nucleatum*: AE009951.530422.531923; *R. gnavus*: AZJF01000012.178214.179732.

| Region | SILVA ID | Start | End |
| --- | --- | --- | --- |
| v2 | *F. nucleatum* | 134 | 389 |
| v2 | *R. gnavus* | 108 | 362 |
| v2 | *B. vulgatus* | 110 | 364 |
| v2 | *B. fragilis* | 108 | 361 |
| v3 | *B. vulgatus* | 330 | 540 |
| v3 | *B. fragilis* | 327 | 537 |
| v4 | *F. nucleatum* | 531 | 818 |
| v4 | *R. gnavus* | 500 | 788 |
| v4 | *B. vulgatus* | 522 | 810 |
| v6v7 | *F. nucleatum* | 944 | 1207 |
| v6v7 | *R. gnavus* | 917 | 1177 |
| v6v7 | *B. vulgatus* | 936 | 1194 |
| v6v7 | *B. fragilis* | 933 | 1193 |

To define the taxonomic structure of the microbiome, we compared three different classifier algorithms which are based on full genome k-mer matching (`Kraken2`), protein-level read alignment (`Kaiju`) or gene specific markers (`MetaPhlAn2`) (Fig. 1a). A common core microbiome structure was observed regardless of the taxonomic classifier method. However, particular deviations in relative abundance were observed between these methods. To estimate the microbiome community structure differences, we performed a PCA of CLR-transformed data, which revealed a clear clustering by the taxonomic classification method (Fig. 2b). Importantly, however, `Kraken2` and `Kaiju` family-level classifications clustered samples in the same order along the second component, which likely reflects consistency in classification despite of the method used.

**Figure 2:**
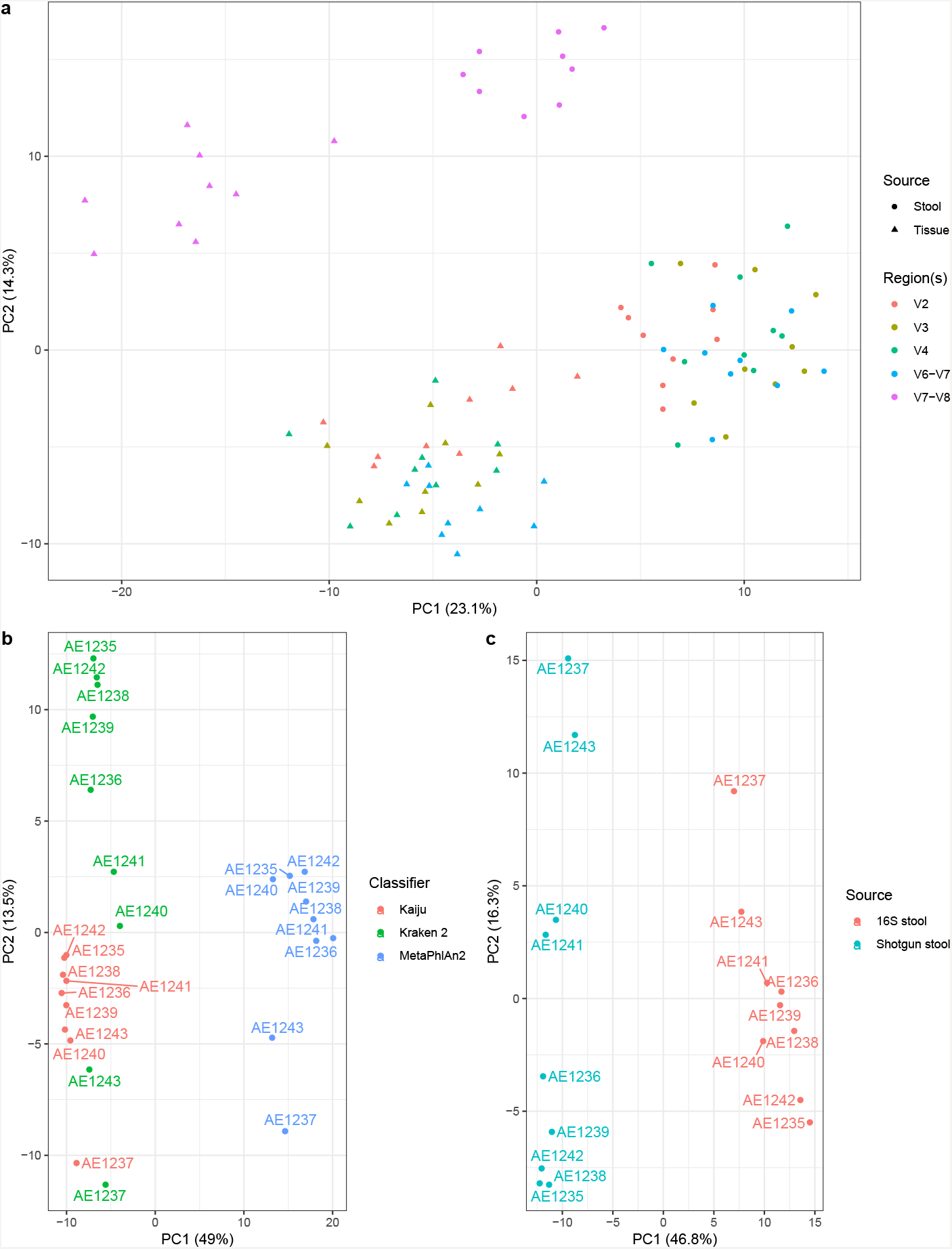
Ordination of samples. Principal components analysis of several datasets after central log ratio transformations of the family-level classifications.16S data, where each sample was separated by region and source material.Shotgun data, classified using Kraken2, Kaiju and MetaPhlAn2. C. 16S data from faeces (only V4 region) and shotgun data (classified using Kraken2).

Likewise, both the variable regions analysed and the source material (faeces or tissue) revealed different distributions of the bacterial taxa (Fig. 1b). Indeed, when analysing CLR-transformed taxonomic profiles, samples clustered mostly by source material (Fig. 2a). Notably, the V7-V8 data showed the largest deviation in principal components from all other variable regions (Fig. 2a).

Lastly, a clear difference in community structure was observed between 16S and shotgun sequences from the same faecal samples (Fig.2c). Regardless, samples were displayed in the same order on the second component, which points to the consistency of this dataset despite technical biases.

We also subsampled original clean reads at lower coverage numbers and computed alpha diversity at different taxonomic and functional classification levels in order to study the sequencing depth necessary to capture microbial diversity (Fig. 3).

**Figure 3:**
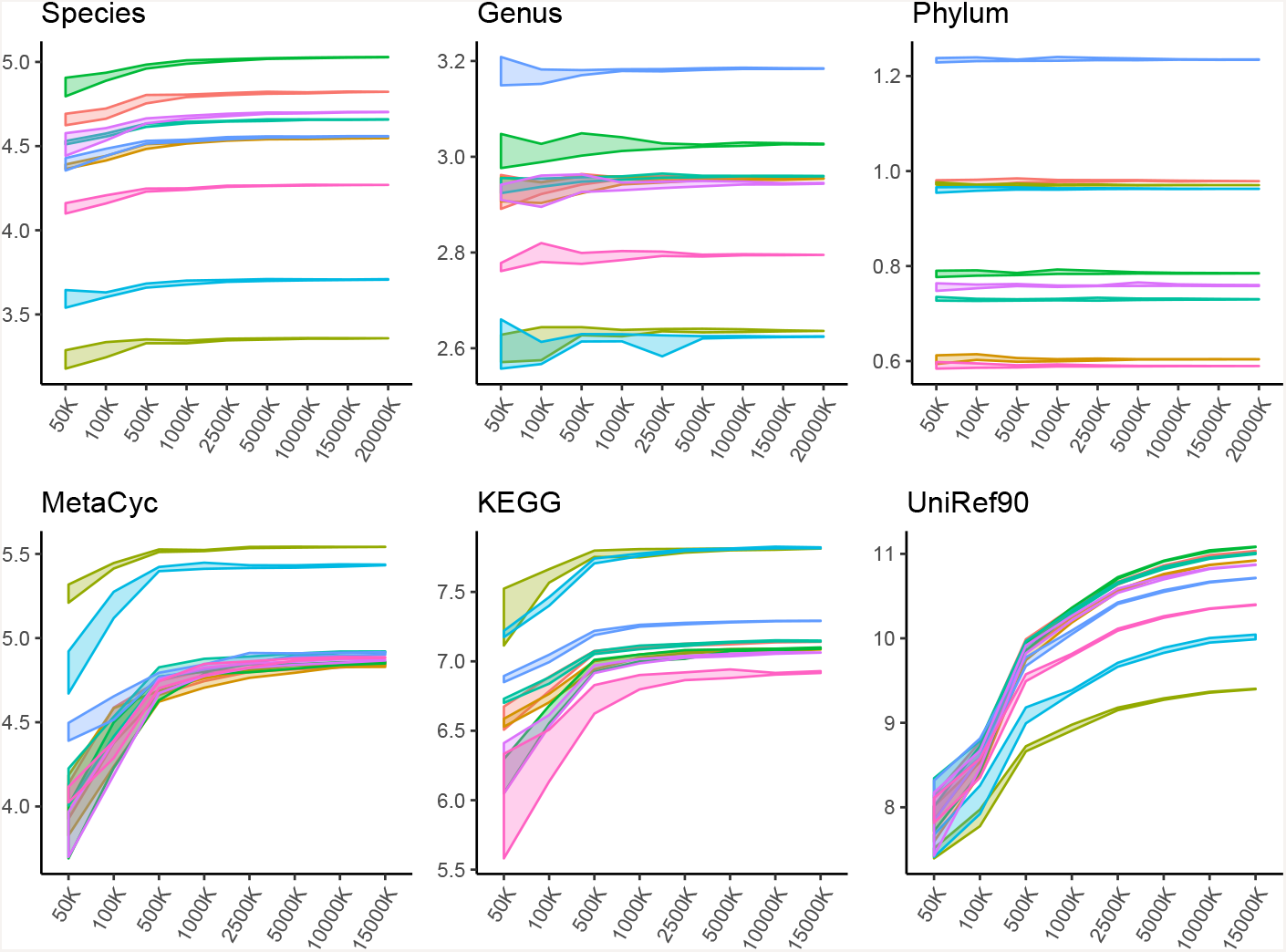
**Shannon alpha diversity index** at different taxonomic (species, genus, phylum) and functional (gene families: UniRef90, functional groups: KEGG orthogroups and metabolic pathways: MetaCyc) by number of read pairs. 5 random samples were created at each level.

These alpha diversity profiles demonstrated a gradual drop in diversity as sequencing coverage decreased. This drop in coverage was more noticeable in features with higher diversity, particularly at species level or when using gene families (UniRef90). Altogether, in the case of species, sequencing coverages as low as 1 million read pairs appeared to capture the taxonomic diversity present in samples, in line with previous findings[50]. Altogether, we demonstrate that our high-coverage dataset from nine participants sustains sufficient sequencing depth to capture the majority of the known bacterial taxa and functional groups present in these samples.

## Usage Notes

For maximum reproducibility purposes, sequencing data was deposited as raw reads. However, human sequencing reads were removed from the dataset prior to uploading in order to prevent participants’ identification. Thus, reads need to be trimmed and, if necessary, deduplicated, before being reutilized.

For 16S data, reads have been uploaded without any manipulation. Hence, reads from different variable regions are present in the same FASTQ file. We suggest researchers to run our reads classification scripts in order to choose variable regions for the analysis. Following that, reads will still need to be quality controlled, either directly or by denoising algorithms such as `DADA2`.

## Acknowledgements

We appreciate the collaboration of all participants who provided epidemiological data and biological samples. We thank all the personnel that were involved in the recruitment process, specially our documentalist Carmen Atencia and our laboratory technician Susana López. This research was financially supported by the Ministry of Science, Innovation and Universities, Government of Spain (grant FPU17/05474). Ministry of Health, Government of Catalonia (grants SLT002/16/00496 and SLT002/16/00398), Spanish Ministry for Economy and Competitivity, Instituto de Salud Carlos III, co-funded by FEDER funds -a way to build Europe– (FIS PI17/00092), Agency for Management of University and Research Grants (AGAUR) of the Catalan Government (grant 2017SGR723). Mireia Obón-Santacana received a post-doctoral fellow from “Fundación Científica de la Asociación Española Contra el Cáncer (AECC)”. We thank CERCA Program, Generalitat de Catalunya for institutional support. None of these agencies had any role in the interpretation of the results or the preparation of this manuscript.

## Author contributions

VP and VM designed and supervised the study. GIS, EG and MOS designed the recruitment protocols. GIS, FRM, AMB and AGR conducted the recruitment and sample collection. MLP and MB contributed to the sample preparation and sequencing protocols. JML conducted the bioinformatics analysis. JML and VP wrote the first draft of the manuscript. All co-authors assisted in the writing of the manuscript and approved the submitted versions.

## References

[1] Maier, L. & Typas, A. Systematically investigating the impact of medication on the gut microbiome. Current Opinion in Microbiology 39, 128–135 (2017).

[2] Maier, L. et al. Extensive impact of non-antibiotic drugs on human gut bacteria. Nature 555, 623–628 (2018).

[3] Goodrich, J. K., Davenport, E. R., Clark, A. G. & Ley, R. E. The Relationship Between the Human Genome and Microbiome Comes into View. Annual Review of Genetics 51, 413–433 (2017).

[4] Parks, D. H. et al. Recovery of nearly 8,000 metagenome-assembled genomes substantially expands the tree of life. Nature Microbiology 2, 1533–1542 (2017).

[5] Almeida, A. et al. A new genomic blueprint of the human gut microbiota. Nature 568, 499–504 (2019).

[6] Pasolli, E. et al. Extensive Unexplored Human Microbiome Diversity Revealed by Over 150,000 Genomes from Metagenomes Spanning Age, Geography, and Lifestyle. Cell 176, 649–662.e20 (2019).

[7] Wirbel, J. et al. Meta-analysis of fecal metagenomes reveals global microbial signatures that are specific for colorectal cancer. Nature Medicine 25 (2019).

[8] Thomas, A. M. et al. Metagenomic analysis of colorectal cancer datasets identifies cross-cohort microbial diagnostic signatures and a link with choline degradation. Nature Medicine In press (2019).

[9] Weisburg, W. G., Barns, S. M., Pelletier, D. A. & Lane, D. J. 16S ribosomal DNA amplification for phylogenetic study. Journal of bacteriology 173, 697–703 (1991).

[10] Yarza, P. et al. Uniting the classification of cultured and uncultured bacteria and archaea using 16S rRNA gene sequences. Nature Reviews Microbiology 12, 635–645 (2014).

[11] Edgar, R. C. Updating the 97% identity threshold for 16S ribosomal RNA OTUs. Bioinformatics 34, 2371–2375 (2018).

[12] Martinez-Porchas, M., Villalpando-Canchola, E., Ortiz Suarez, L. E. & Vargas-Albores, F. How conserved are the conserved 16S-rRNA regions? PeerJ 5, e3036 (2017).

[13] Yang, B., Wang, Y. & Qian, P. Y. Sensitivity and correlation of hypervariable regions in 16S rRNA genes in phylogenetic analysis. BMC Bioinformatics 17, 1–8 (2016).

[14] Barb, J. J. et al. Development of an Analysis Pipeline Characterizing Multiple Hypervariable Regions of 16S rRNA Using Mock Samples. PLoS ONE 11, 1–18 (2016).

[15] D’Amore, R. et al. A comprehensive benchmarking study of protocols and sequencing platforms for 16S rRNA community profiling. BMC Genomics 17 (2016).

[16] Lindgreen, S., Adair, K. L. & Gardner, P. P. An evaluation of the accuracy and speed of metagenome analysis tools. Scientific Reports 6, 1–14 (2016).

[17] McIntyre, A. B. et al. Comprehensive benchmarking and ensemble approaches for metagenomic classifiers. Genome Biology 18, 1–19 (2017).

[18] Truong, D. T., Tett, A., Pasolli, E., Huttenhower, C. & Segata, N. Microbial strain-level population structure and genetic diversity from metagenomes. Genome Research 27, 626–638 (2017). arXiv:1312.0570v2.

[19] van der Walt, A. J. et al. Assembling metagenomes, one community at a time. BMC Genomics 18, 1–13 (2017).

[20] Vincent, A. T., Derome, N., Boyle, B., Culley, A. I. & Charette, S. J. Next-generation sequencing (NGS) in the microbiological world: How to make the most of your money. Journal of Microbiological Methods 138, 60–71 (2017).

[21] Walsh, A. M. et al. Species classifier choice is a key consideration when analysing low-complexity food microbiome data. Microbiome 6, 50 (2018).

[22] Breitwieser, F. P., Lu, J. & Salzberg, S. L. A review of methods and databases for metagenomic classification and assembly. Briefings in Bioinformatics (2017).

[23] Clooney, A. G. et al. Comparing apples and oranges?: Next generation sequencing and its impact on microbiome analysis. PLoS ONE 11, 1–16 (2016).

[24] Jovel, J. et al. Characterization of the gut microbiome using 16S or shotgun metagenomics. Frontiers in Microbiology 7, 1–17 (2016).

[25] Laudadio, I. et al. Quantitative Assessment of Shotgun Metagenomics and 16S rDNA Amplicon Sequencing in the Study of Human Gut Microbiome. OMICS: A Journal of Integrative Biology 22, 248–254 (2018).

[26] Jones, R. B. et al. Inter-niche and inter-individual variation in gut microbial community assessment using stool, rectal swab, and mucosal samples. Scientific Reports 8, 1–12 (2018).

[27] JoanML/colonbiome-pilot. https://gitlab.com/JoanML/colonbiome-pilot.

[28] Peris, M. et al. Lessons learnt from a population-based pilot programme for colorectal cancer screening in Catalonia (Spain). Journal of Medical Screening 14, 81–86 (2007).

[29] Binefa, G. et al. Colorectal Cancer Screening Programme in Spain: Results of Key Performance Indicators after Five Rounds (2000-2012). Scientific Reports 6, 1–10 (2016).

[30] Atkin, W. S. et al. European guidelines for quality assurance in colorectal cancer screening and diagnosisFirst Edition Colonoscopic surveillance following adenoma removal. Endoscopy 44, 151–163 (2012).

[31] Gloor, G. B., Macklaim, J. M., Pawlowsky-Glahn, V. & Egozcue, J. J. Microbiome Datasets Are Compositional: And This Is Not Optional. Frontiers in Microbiology 8 (2017).

[32] Langmead, B. & Salzberg, S. L. Fast gapped-read alignment with Bowtie 2. Nature Methods 9, 357–359 (2012).

[33] Li, H. et al. The Sequence Alignment/Map format and SAMtools. Bioinformatics (Oxford, England) 25, 2078–9 (2009).

[34] FASTQC. https://www.bioinformatics.babraham.ac.uk/projects/fastqc/.

[35] BBTools. https://jgi.doe.gov/data-and-tools/bbtools/.

[36] Wood, D. E. & Salzberg, S. L. Kraken: Ultrafast metagenomic sequence classification using exact alignments. Genome Biology 15 (2014).

[37] Lu, J., Breitwieser, F. P., Thielen, P. & Salzberg, S. L. Bracken: estimating species abundance in metagenomics data. PeerJ Computer Science 3, e104 (2017).

[38] Truong, D. T. et al. MetaPhlAn2 for enhanced metagenomic taxonomic profiling. Nature Methods 12, 902–903 (2015).

[39] Menzel, P., Ng, K. L. & Krogh, A. Fast and sensitive taxonomic classification for metagenomics with Kaiju. Nature Communications 7, 1–9 (2016).

[40] Franzosa, E. A. et al. Functionally profiling metagenomes and metatranscriptomes at species-level resolution. Nature methods accepted (2018).

[41] Nurk, S., Meleshko, D., Korobeynikov, A. & Pevzner, P. A. metaSPAdes: a new versatile metagenomic assembler. Genome research 27, 824–834 (2017).

[42] Kang, D. et al. MetaBAT 2: an adaptive binning algorithm for robust and efficient genome reconstruction from metagenome assemblies. PeerJ Preprints 27522v1 (2019).

[43] Parks, D. H., Imelfort, M., Skennerton, C. T., Hugenholtz, P. & Tyson, G. W. CheckM: assessing the quality of microbial genomes recovered from isolates, single cells, and metagenomes. Genome research 25, 1043–55 (2015).

[44] Segata, N., Börnigen, D., Morgan, X. C. & Huttenhower, C. PhyloPhlAn is a new method for improved phylogenetic and taxonomic placement of microbes. Nature Communications 4 (2013).

[45] Callahan, B. J. et al. DADA2: High-resolution sample inference from Illumina amplicon data. Nature Methods 13, 581–583 (2016). 15334406.

[46] Murali, A., Bhargava, A. & Wright, E. S. IDTAXA: A novel approach for accurate taxonomic classification of microbiome sequences. Microbiome 6, 1–14 (2018).

[47] Oksanen, J. et al. vegan: Community Ecology Package (2019). URL https://CRAN.R-project.org/package=vegan. R package version 2.5-5.

[48] Palarea-Albaladejo, J. & Martín-Fernández, J. A. zCompositions — R package for multivariate imputation of left-censored data under a compositional approach. Chemometrics and Intelligent Laboratory Systems 143, 85–96 (2015).

[49] CoDaSeq. https://github.com/ggloor/CoDaSeq.

[50] Hillmann, B. et al. Evaluating the Information Content of Shallow Shotgun Metagenomics. mSystems 3, 1–12 (2018).

